# Insular Celtic population structure and genomic footprints of migration

**DOI:** 10.1101/230797

**Authors:** Ross P Byrne, Rui Martiniano, Lara M Cassidy, Matthew Carrigan, Garrett Hellenthal, Orla Hardiman, Daniel G Bradley, Russell L McLaughlin

## Abstract

Previous studies of the genetic landscape of Ireland have suggested homogeneity, with population substructure undetectable using single-marker methods. Here we have harnessed the haplotype-based method fineSTRUCTURE in an Irish genome-wide SNP dataset, identifying 23 discrete genetic clusters which segregate with geographical provenance. Cluster diversity is pronounced in the west of Ireland but reduced in the east where older structure has been eroded by historical migrations. Accordingly, when populations from the neighbouring island of Britain are included, a west-east cline of Celtic-British ancestry is revealed along with a particularly striking correlation between haplotypes and geography across both islands. A strong relationship is revealed between subsets of Northern Irish and Scottish populations, where discordant genetic and geographic affinities reflect major migrations in recent centuries. Additionally, Irish genetic proximity of all Scottish samples likely reflects older strata of communication across the narrowest inter-island crossing. Using GLOBETROTTER we detected Irish admixture signals from Britain and Europe and estimated dates for events consistent with the historical migrations of the Norse-Vikings, the Anglo-Normans and the British Plantations. The influence of the former is greater than previously estimated from Y chromosome haplotypes. In all, we paint a new picture of the genetic landscape of Ireland, revealing structure which should be considered in the design of studies examining rare genetic variation and its association with traits.

**Author summary:** A recent genetic study of the UK (People of the British Isles; PoBI) expanded our understanding of population history of the islands, using newly-developed, powerful techniques that harness the rich information embedded in chunks of genetic code called haplotypes. These methods revealed subtle regional diversity across the UK, and, using genetic data alone, timed key migration events into southeast England and Orkney. We have extended these methods to Ireland, identifying regional differences in genetics across the island that adhere to geography at a resolution not previously reported. Our study reveals relative western diversity and eastern homogeneity in Ireland owing to a history of settlement concentrated on the east coast and longstanding Celtic diversity in the west. We show that Irish Celtic diversity enriches the findings of PoBI; haplotypes mirror geography across Britain and Ireland, with relic Celtic populations contributing greatly to haplotypic diversity. Finally, we used genetic information to date migrations into Ireland from Europe and Britain consistent with historical records of Viking and Norman invasions, demonstrating the signatures of these migrations the on modern Irish genome. Our findings demonstrate that genetic structure exists in even small isolated populations, which has important implications for population-based genetic association studies.

## Introduction

Situated at the northwestern edge of Europe, Ireland is the continent’s third largest island, with a modern-day population of approximately 6.4 million. The island is politically partitioned into the Republic of Ireland and Northern Ireland, with the latter forming part of the United Kingdom (UK) alongside the neighbouring island of Britain. Alternative divisions separate Ireland into four provinces reflecting early historical divisions: Ulster to the north, including Northern Ireland; Leinster (east); Munster (south) and Connacht (west). Humans have continuously inhabited Ireland for around 10,000 years [1], though it is not until after the demographic upheavals of the Early Bronze Age (circa 2200 BCE), that strong genetic continuity between ancient and modern Irish populations is observed [2]. Linguistically, the island’s earliest attested language forms part of the Insular Celtic family, specifically the Gaelic branch, whose historic range also extended to include many regions of Scotland, via maritime connections with Ulster [3,[4]. A second branch of Insular Celtic, the Brittonic languages, had been spoken across much of Britain up until the introduction of Anglo-Saxon in the 5th and 6th centuries, by which time they were diversifying into Cornish, Welsh and Cumbric dialects [5].

Since the establishment of written history, numerous settlements and invasions of Ireland from the neighbouring island of Britain and continental Europe have been recorded. This includes Norse-Vikings (9th-12th century), especially in east Leinster, and Anglo-Normans (12th-14th century), who invaded through Wexford in the southeast and established English rule mainly from an area later called the Pale in northeast Leinster [6]. There has also been continuous movement of people from Britain, in particular during the 16-17th century Plantation periods during which Gaelic and Norman lands were systematically colonized by English and Scottish settlers. These events had a particularly enduring impact in Ulster in comparison with other planted regions such as Munster. As with the previous Norman invasion, the less fertile west of the country (Connacht) remained largely untouched during this period.

The genetic contributions of these migratory events cannot be considered mutually independent, given that they derive from either related Germanic populations (such as the Vikings and their purported Norman descendants) or from other Celtic populations inhabiting Britain, which had themselves been subjected to mass Germanic influx from Anglo-Saxon migrations and later Viking and Norman invasions [7]. Moreover, each movement of people originated from northern Europe, a region which had witnessed a mass homogenizing of genetic variation during the migrations of the Early Bronze Age, possibly linked to Indo-European language spread. [8,[9]. However, each event had a geographic and temporal focal point on the island, which may be detectable in local population structure.

Previous genome-wide surveys have detected little to no structure in Ireland using methods such as principal component analysis (PCA) on independent markers, concluding that the Irish population is genetically homogenous [10]. However, runs of homozygosity are relatively long and frequent in Ireland [10] and correlate negatively with population density and diversity of grandparental origins [11], suggesting that low ancestral mobility may have preserved regional genetic legacies within Ireland, which may be detectable in modern genomes as local population structure embedded within haplotypes. This is further supported by the restricted regional distributions of certain Y chromosome haplotypes [12,[13].

The haplotype-based methods ChromoPainter and fineSTRUCTURE [14] were recently used to uncover hidden genetic structure among the people of modern Britain [7]. These approaches exploit the rich information available within haplotypes (usually statistically phased) to identify clusters of genetically distinct individuals with a resolution that could not be attained using single-marker methods. In doing so, the People of the British Isles (PoBI) study was able to identify discrete genetic clusters of individuals that strongly segregate with geographical regions within Britain, though notably, structure was undetectable across a large southeastern portion of the island. However, although this study sampled over 2,000 individuals, only 44 were from Northern Ireland with none from the remainder of the island. Ireland was also excluded from admixture and ancestry analyses due to the confounding effects of the island acting as “a source and a sink for ancestry from the UK”. With this focus on a single island, the PoBI study has an obvious limit, despite its title.

Here, we have used the methods of the PoBI study to explore fine-grained Irish population substructure. We first investigate Ireland on its own, then we consider the genetic substructure observed on the island in the context of Britain and continental Europe. Using modern individuals from these two sources as surrogates for historical populations, we apply the GLOBETROTTER model to infer admixture events into Ireland and we consider these in the context of historically recorded invasions and migrations. Our inclusion of Irish data with previously-published data from Britain presents a more complete representation of genetic ancestry in the contemporary populations of the British Isles, providing a comprehensive population genetic perspective of the peopling of these islands.

## Results and discussion

### Celtic population structure in Ireland

We used ChromoPainter [14] to identify haplotypic similarities within a genome-wide single nucleotide polymorphism (SNP) dataset of individuals from the Republic of Ireland and Northern Ireland (n=1,035, including 44 from the PoBI study), in which local geographic origin was known for a subset (n=588). Clustering the resulting coancestry matrix using fineSTRUCTURE identified 23 clusters, demonstrating local population structure within Ireland to a level not previously reported, with apparent geographical, sociopolitical and ancestral correlates (Fig 1). All clusters were robustly defined, with total variation distance (TVD) p-values less than 0.001 (S1 and S2 Table). We projected the ChromoPainter coancestry matrix in lower-dimensional space using principal component analysis (PCA) and, to ease interpretation and for visual brevity with labels, we defined 9 cluster groups that formed higher order clades in the fineSTRUCTURE dendrogram, overlapped in PC space and were sampled from geographically contiguous regions. These cluster groups also showed robust definition by TVD analysis (S3 Table and S4 Table), suggesting they are meaningful. ChromoPainter PCA revealed a tight relationship between haplotypic similarity and geographical proximity, with principal component (PC) 1 roughly describing a north to south cline and PC2 largely describing an east to west cline (Fig 1B).

**Fig 1.**
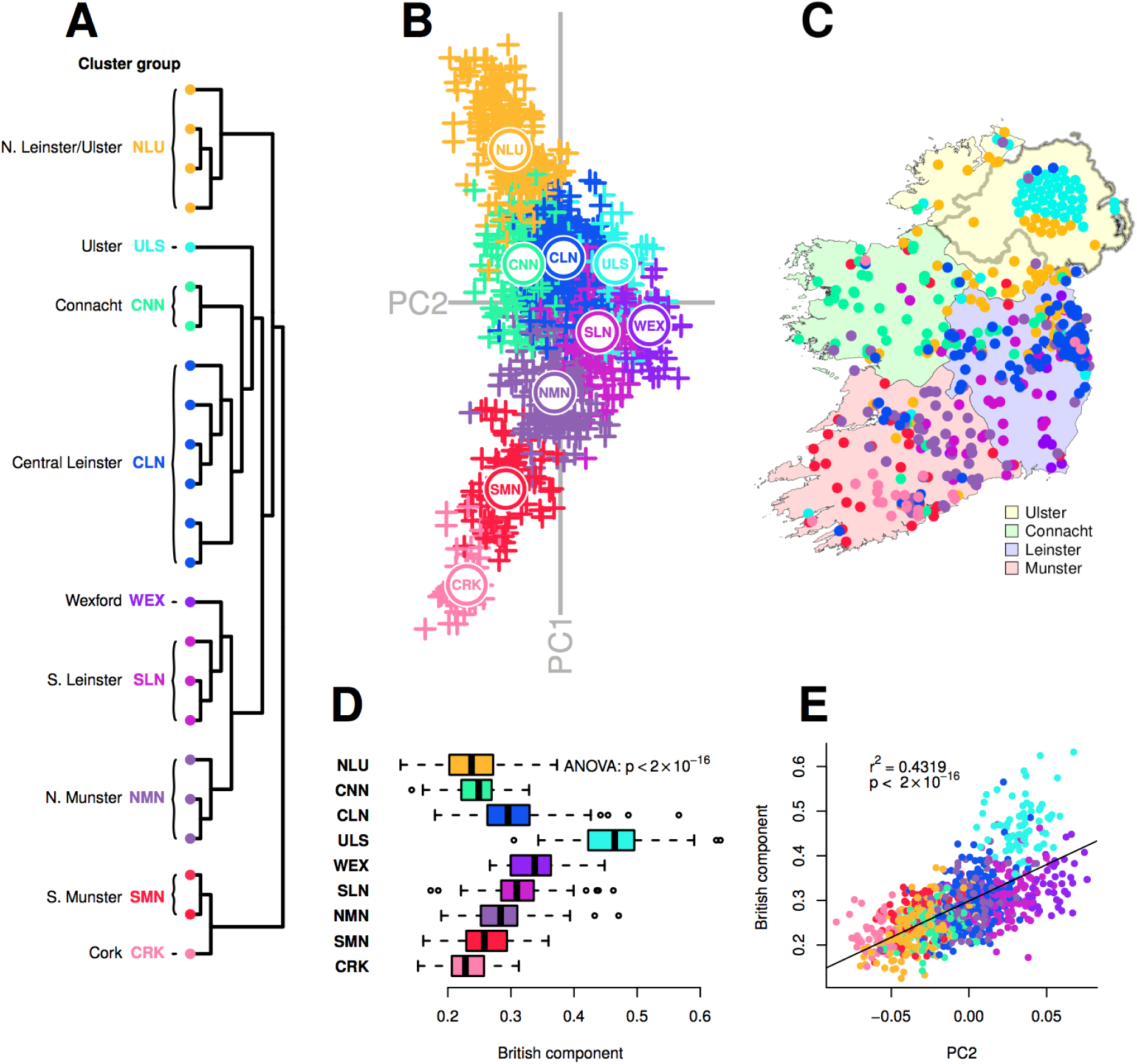
Fine-grained population structure in Ireland. **(A)** fineSTRUCTURE clustering dendrogram for 1,035 Irish individuals. Twenty-three clusters are defined, which are combined into cluster groups for clusters that are neighbouring in the dendrogram, overlapping in principal component space (B) and sampled from regions that are geographically contiguous. Details for each cluster in the dendrogram are provided in S1 Fig. **(B)** Principal components analysis (PCA) of haplotypic similarity, based on ChromoPainter coancestry matrix for Irish individuals. Points are coloured according to cluster groups defined in (A); the median location of each cluster group is plotted. **(C)** Map of Ireland showing the sampling location for a subset of 588 individuals analysed in (A) and (B), coloured by cluster group. Points have been randomly jittered within a radius of 5 km to preserve anonymity. Precise sampling location for 44 Northern Irish individuals from the People of the British Isles dataset was unknown; these individuals are plotted geometrically in a circle. **(D)** “British admixture component” (ADMIXTURE estimates; k=2) for Irish cluster groups. This component has the largest contribution in ancient Anglo-Saxons and the SEE cluster. **(E)** Linear regression of principal component 2 (B) versus British admixture component (r^2^ = 0.43; p < 2×10^−16^). Points are coloured by cluster group. (Standard error for ADMIXTURE point estimates presented in S11 Fig.)

At a high level, both ChromoPainter PCA and fineSTRUCTURE clustering loosely separated the historical provinces of Ireland (Ulster, Leinster, Munster and Connacht) suggesting that these socially constructed territories may have had an impact on genetic structure within Ireland which is deeply embedded in time. Careful inspection of the tree ordering and the PCA revealed more nuanced relations between the provinces; for example south Leinster clusters share more haplotypes with those from north Munster than with their central and north Leinster counterparts. The geographical distribution of this deep subdivision of Leinster resembles pre-Norman territorial boundaries which divided Ireland into fifths (*cúige*), with north Leinster a kingdom of its own known as Meath (*Mide*) [15]. However interpreted, the firm implication of the observed clustering is that despite its previously reported homogeneity, the modern Irish population exhibits genetic structure that is subtly but detectably affected by ancestral population structure conferred by geographical distance and, possibly, ancestral social structure.

ChromoPainter PC1 demonstrated high diversity amongst clusters from the west coast, which may be attributed to longstanding residual ancient (possibly Celtic) structure in regions largely unaffected by historical migration. Alternatively, genetic clusters may also have diverged as a consequence of differential influence from outside populations. This diversity between western genetic clusters cannot be explained in terms of geographic distance alone. South Munster (SMN) and Cork (CRK) clusters branch off first in the fineSTRUCTURE tree and show distinct separation from their neighbouring north Munster clusters (NMN), indicating that south Munster’s haplotypic makeup is more distinct from its neighbouring regions and the remaining regions than any other cluster. TVD analysis supports this observation (S1 Table and S3 Table), with the Cork cluster in particular showing strong differentiation from other clusters. This may reflect the persistent isolating effects of the mountain ranges surrounding the south Munster counties of Cork and Kerry, restricting gene flow with the rest of Ireland and preserving older structure.

In contrast to the west of Ireland, eastern individuals exhibited relative homogeneity; a similar pattern was observed in the PoBI study [7], in which all samples in a large region in southeast England formed a single indivisible cluster of genetically similar individuals comprising almost half the dataset. However, while east coast clusters in Ireland are the largest and demonstrate strong cluster integrity, the largest of these (the Central Leinster Cluster, CLN) comprises roughly a fifth of our dataset (S1 Fig), hence they are dwarfed proportionally in both number and geographical extent by the southeast England cluster, suggesting that deeper structure persists in eastern Ireland than in southeast England. The overall pattern of western diversity and eastern homogeneity in Ireland may be explained by increased gene flow and migration into and across the east coast of Ireland from geographically proximal regions, the closest of which is the neighbouring island of Britain.

To explore this, we estimated the extent of admixture per individual in the Irish dataset from Britain, using samples from the PoBI dataset as a reference [7], along with eighteen ancient British individuals from the Iron Age, Roman and Anglo-Saxon periods in northeast and southeast England [16,[17]. Using an unsupervised ADMIXTURE analysis [18], we observed that one of the ADMIXTURE clusters (k=2) comprises the totality of ancestry of several Anglo-Saxon individuals and forms the largest proportion in British groups, with varying representation across Irish clusters (S8 Fig). For simplicity we will call this the British component, which was among the lowest for individuals falling in Irish west coast fineSTRUCTURE clusters, including the south Munster and Cork cluster groups (Fig 1D), supporting the interpretation that these regions differ in terms of restricted haplotypic contribution from Britain. Analysis of variance of the British admixture component in cluster groups showed a significant difference (p < 2×10^−16^), indicating a role for British Anglo-Saxon admixture in distinguishing clusters, and ChromoPainter PC2 was correlated with the British component (p < 2×10^−16^), explaining approximately 43% of the variance. PC2 therefore captures an east to west Anglo-Celtic cline in Irish ancestry. This may explain the relative eastern homogeneity observed in Ireland, which could be a result of the greater English influence in Leinster and the Pale during the period of British rule in Ireland following the Norman invasion, or simply geographic proximity of the Irish east coast to Britain. Notably, the Ulster cluster group harboured an exceptionally large proportion of the British component (Fig 1D and 1E), undoubtedly reflecting the strong influence of the Ulster Plantations in the 17th century and its residual effect on the ethnically British population that has remained.

### The genetic structure of the British Isles

The genetic substructure observed in Ireland is consistent with long term geographic diversification of Celtic populations and the continuity shown between modern and Early Bronze Age Irish people [2]. However, this diversity is weaker on the east coast in a manner that correlates with British admixture, suggesting a role for recent migrations in eroding this structure. We therefore further investigated the relationship between Ireland and Britain by generating a ChromoPainter coancestry matrix for all Irish and PoBI data combined (n=3,008). Clustering with fineSTRUCTURE revealed 50 distinct clusters that segregated geographically, both on a cohort-wide and local level (Fig 2). Projecting this coancestry matrix in PC space revealed a striking concordance between haplotypes and geography (sampling regions were defined using Nomenclature of Territorial Units for Statistics 2010; [19]) for ChromoPainter PCs 1 and 4, reminiscent of previous observations for single marker-based summaries of genetic variation within European populations [20].

**Fig 2.**
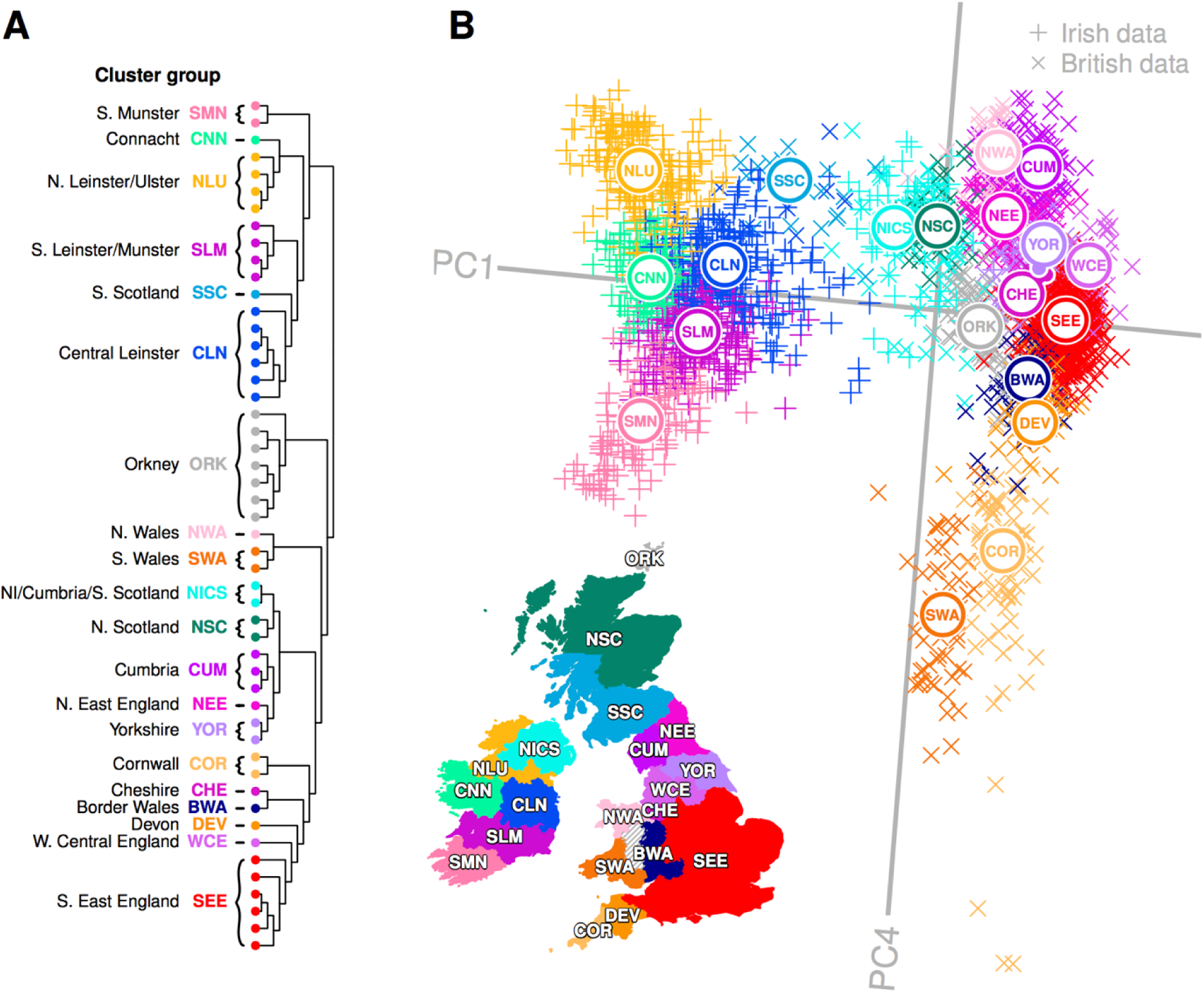
Genes mirror geography in the British Isles. (A) fineSTRUCTURE clustering dendrogram for combined Irish and British data. Data principally split into Irish and British groups before subdividing into a total of 50 distinct clusters, which are combined into cluster groups for clusters that formed clades in the dendrogram, overlapped in principal component space (B) and were sampled from regions that are geographically contiguous. Names and labels follow the geographical provenance for the majority of data within the cluster group. Details for each cluster in the dendrogram are provided in S2 Fig. (B) Principal component analysis (PCA) of haplotypic similarity based on the ChromoPainter coancestry matrix, coloured by cluster group with their median locations labelled. We have chosen to present PC1 versus PC4 here as these components capture new information regarding correlation between haplotypic variation across Britain and Ireland and geography, while PC2 and PC3 (Fig 4) capture previously reported splitting for Orkney and Wales from Britain [7]. A map of Ireland and Britain is shown for comparison, coloured by sampling regions for cluster groups, the boundaries of which are defined by the Nomenclature of Territorial Units for Statistics (NUTS 2010), with some regions combined. Sampling regions are coloured by the cluster group with the majority presence in the sampling region; some sampling regions have significant minority cluster group representations as well, for example the Northern Ireland sampling region (UKN0; NUTS 2010) is majorly explained by the NICS cluster group but also has significant representation from the NLU cluster group. The PCA plot has been rotated clockwise by 5 degrees to highlight its similarity with the geographical map of the Ireland and Britain. NI, Northern Ireland; PC, principal component. Cluster groups that share names with groups from Fig 1 (NLU; SMN; CLN; CNN) have an average of 80% of their samples shared with the initial cluster groups. © EuroGeographics for the map and administrative boundaries, note some boundaries have been subsumed or modified to better reflect sampling regions.

The principal split in the combined Irish and British data defined two genetic islands, both in the fineSTRUCTURE tree and in ChromoPainter PC1 (Fig 2). This distinction between Irish and British genetic data was particularly pronounced when we applied t-distributed stochastic neighbour embedding (t-SNE) [21] to the ChromoPainter coancestry matrix (Fig 3). t-SNE is a nonlinear dimensionality reduction method that attempts to provide an optimal low-dimensional embedding of data by preserving both local and global structure, placing similar points close to each other and dissimilar points far apart. In principle, a two-dimensional t-SNE plot can therefore summarize more of the overall differences between groups than those described by any two principal components, although the relative group sizes, positions and distances on the plot are less straightforward to interpret. Applying t-SNE to the Irish and British coancestry matrix captured the salient structure described by PCA, and particularly validates that observed in the plot of ChromoPainter PC1 vs PC4. This clearly distinguishes the two islands, discerns their north-south and west-east genetic structure and places Orkney and north/south Wales, whose variation is captured in ChromoPainter PCs 2 and 3 respectively (Fig 4), as independent entities from the bulk of the British data.

**Fig 3.**
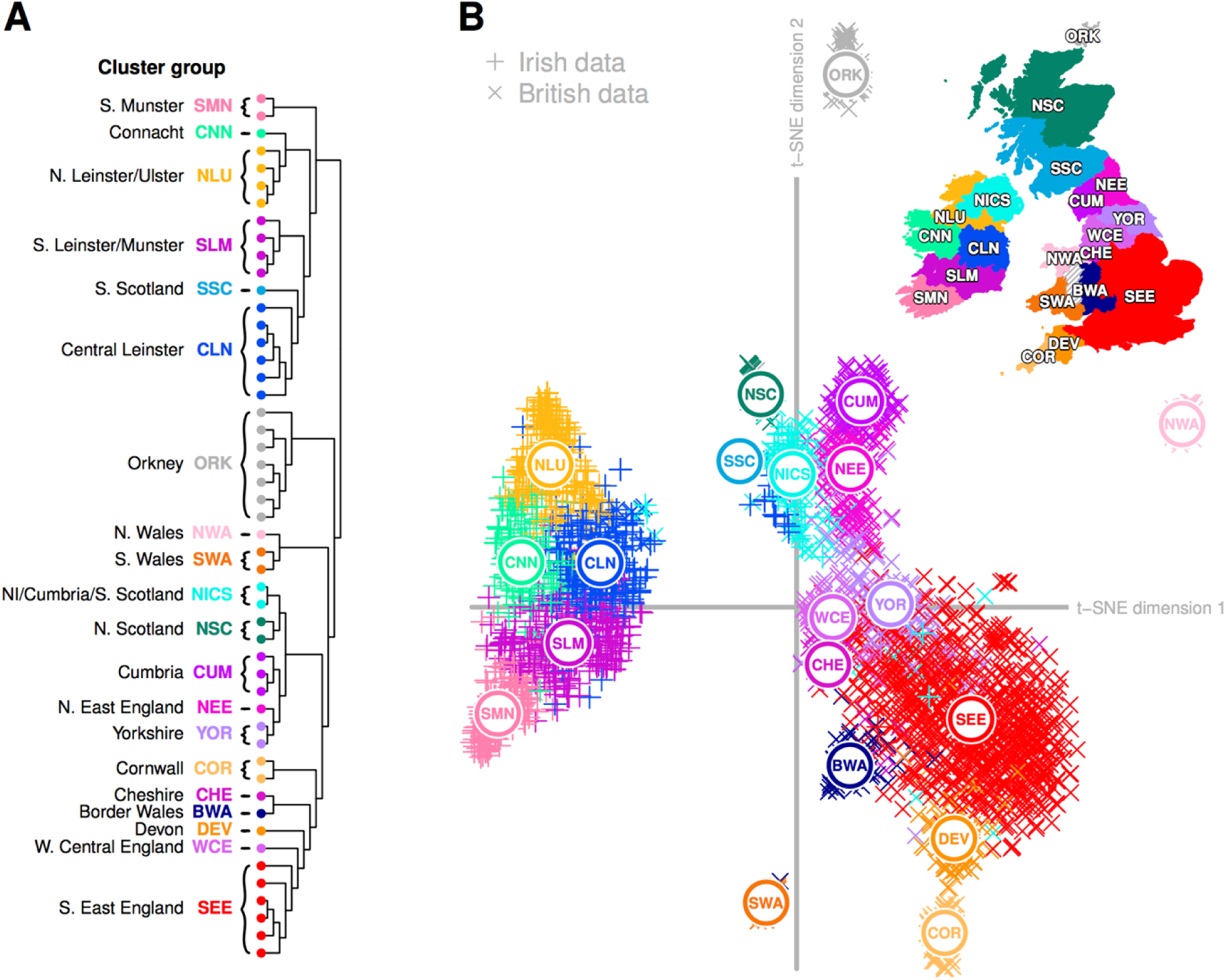
t-distributed stochastic neighbour embedding (t-SNE) of Irish and British coancestry matrix. **(A)** fineSTRUCTURE dendrogram with clusters and cluster groups defined as in Fig 2. **(B)** Two-dimensional t-SNE embedding of ChromoPainter coancestry matrix, with median locations for cluster groups plotted. As t-SNE is a stochastic method, different runs produce different solutions to the 2-dimensional embedding; shown here is a typical result. t-SNE performed significantly better with the ChromoPainter coancestry matrix than with Hamming distances (identity-by-state) computed over single SNP markers (S9 Fig). © EuroGeographics for the map and administrative boundaries, note some boundaries have been subsumed or modified to better reflect sampling regions.

**Fig 4:**
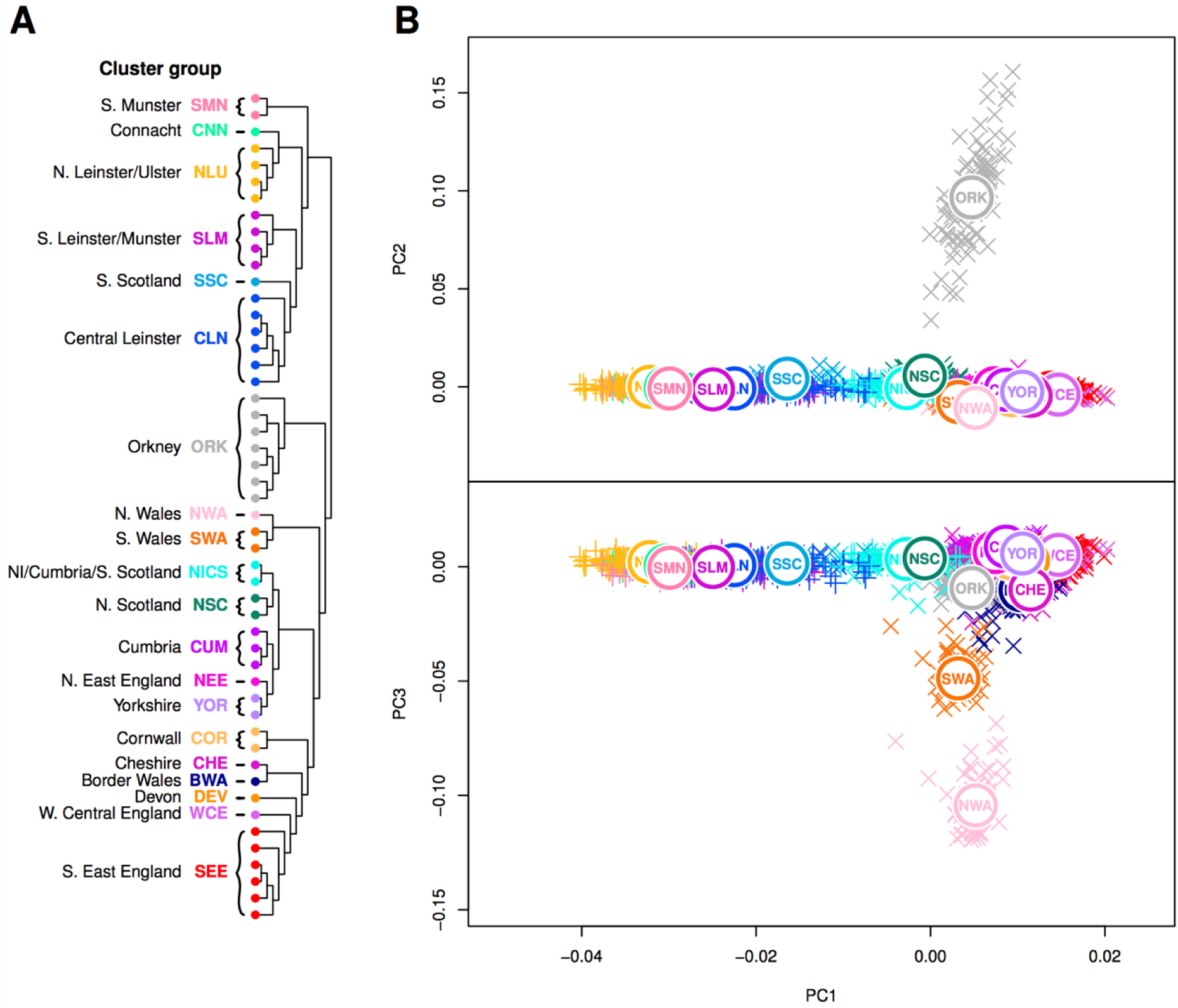
Principal components 2 and 3 of combined Irish and British coancestry matrix. **(A)** fineSTRUCTURE clustering dendrogram for combined Irish and British data, with cluster groups defined as in Fig 2. Immediately following the principal inter-island split, Orkney and Wales branch in sequence, consistent with previous observations. **(B)** Principal component analysis (PCA) of haplotypic similarity based on the ChromoPainter coancestry matrix, coloured by cluster group with their median locations labelled. PC2 captures an Orkney split, while PC3 captures a Welsh split.

As observed in Fig 1, ChromoPainter PCA in Ireland and Britain (Fig 2) demonstrates eastern homogeneity for each island and relative diversity on the west coast. The southeast England (SEE) cluster group anchors PC4, centred at zero, representing a group with predominantly Anglo-Saxon-like ancestry (S8 Fig). Clusters representing Celtic populations harbouring less Anglo-Saxon influence separate out above and below SEE on PC4. Notably, northern Irish clusters (NLU), Scottish (NISC, SSC and NSC), Cumbria (CUM) and North Wales (NWA) all separate out at a mutually similar level, representing northern Celtic populations. The southern Celtic populations Cornwall (COR), south Wales (SWA) and south Munster (SMN) also separate out on similar levels, indicating some shared haplotypic variation between geographically proximate Celtic populations across both Islands. It is notable that after the split of the ancestrally divergent Orkney, successive ChromoPainter PCs describe diversity in British populations where “Anglo-saxonization” was repelled [22]. PC3 is dominated by Welsh variation, while PC4 in turn splits North and South Wales significantly, placing south Wales adjacent to Cornwall and north Wales at the other extreme with Cumbria, all enclaves where Brittonic languages persisted.

Scotland is another region of Britain which successfully retained its Celtic language, however in contrast to Welsh and Cornish clusters, the majority of Scottish variation is described by ChromoPainter PC1. The three definable Scottish groups do not drive any further components of variation (up to ChromoPainter PC7 considered) and fall away from the bulk of British variation on PC1, towards Irish clusters. This is most strikingly observed for the southern Scottish cluster (SSC) which fell amongst Irish branches in the fineSTRUCTURE tree, overlapping with samples from the north of Ireland in ChromoPainter PC space (Fig 2 and Fig 5). In an interesting symmetry, many Northern Irish samples clustered strongly with southern Scottish and northern English samples, defining the Northern Irish/Cumbrian/Scottish (NICS) cluster group. More generally, by modelling Irish genomes as a linear mixture of haplotypes from British clusters, we found that Scottish and northern English samples donated more haplotypes to clusters in the north of Ireland than to the south, reflecting an overall correlation between Scottish/north English contribution and ChromoPainter PC1 position in Fig 1 (Linear regression: p < 2×10^−16^, r^2^ = 0.24).

**Fig 5.**
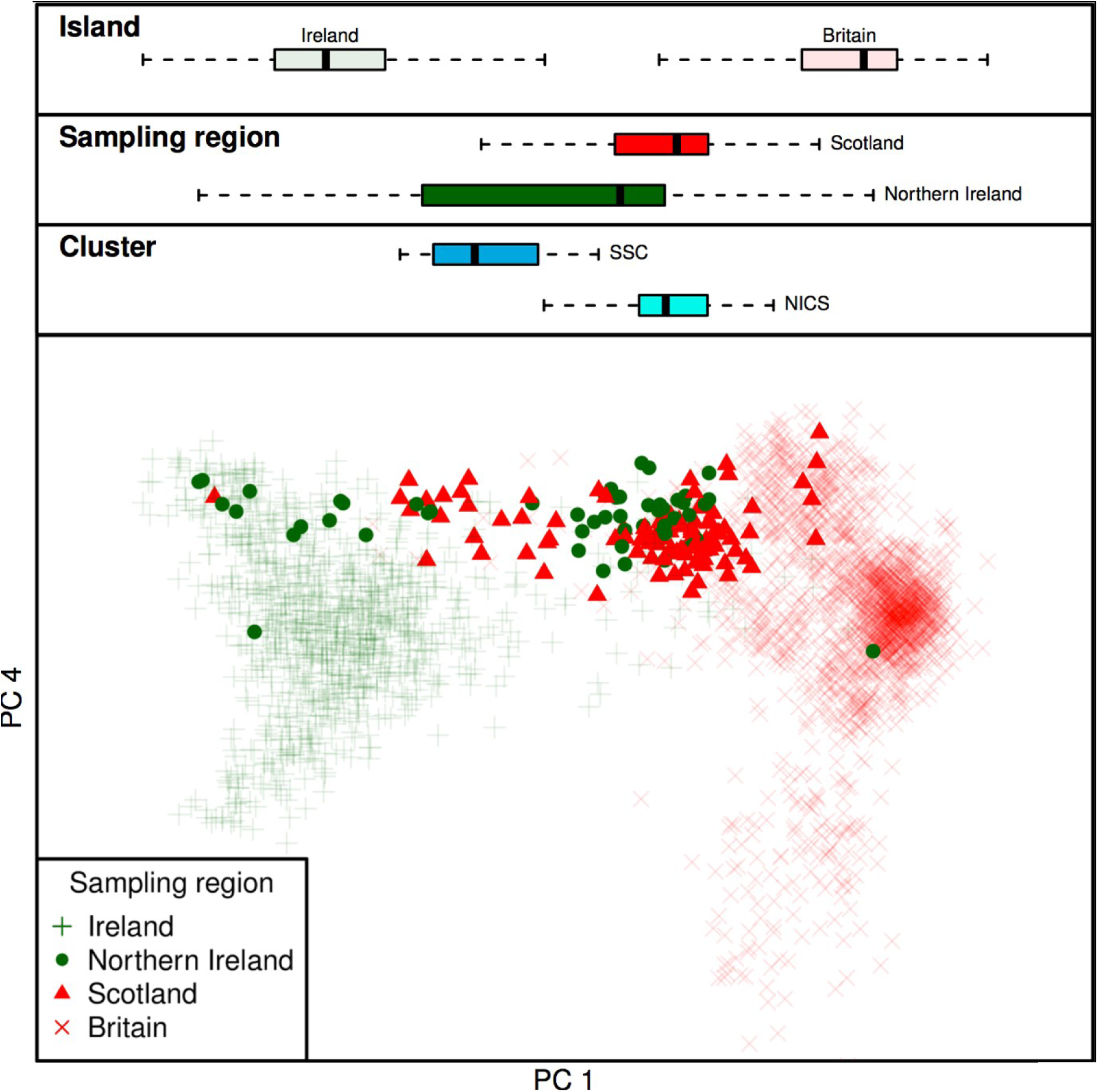
Inter-island exchange of haplotypes between the north of Ireland and northern Britain. The boxplots show the distribution of individuals on principal component (PC) 1 for each island and for specific sampling regions (Scotland/Northern Ireland) and cluster groups (SSC and NICS; see Fig 2). A substantial proportion of Northern Irish individuals fall within the expected range for Scottish individuals in PC space and *vice versa*. This exchange is particularly pronounced for Northern Irish and Scottish individuals that fall within the NICS and SSC cluster groups (Fig 2), respectively.

North to south variation in Ireland and Britain are therefore not independent, reflecting major gene flow between the north of Ireland and Scotland (Fig 5) which resonates with three layers of historical contacts. First, the presence of individuals with strong Irish affinity among the third generation PoBI Scottish sample can be plausibly attributed to major economic migration from Ireland in the 19th and 20th centuries [6]. Second, the large proportion of Northern Irish who retain genomes indistinguishable from those sampled in Scotland accords with the major settlements (including the Ulster Plantation) of mainly Scottish farmers following the 16th Century Elizabethan conquest of Ireland which led to these forming the majority of the Ulster population. Third, the suspected Irish colonisation of Scotland through the *Dál Riata* maritime kingdom, which expanded across Ulster and the west coast of Scotland in the 6th and 7th centuries, linked to the introduction and spread of Gaelic languages [3]. Such a migratory event could work to homogenise older layers of Scottish population structure, in a similar manner as noted on the east coasts of Britain and Ireland. Earlier communications and movements across the Irish Sea are also likely, which at its narrowest point separates Ireland from Scotland by approximately 20 km.

### Genomic footprints of migration into Ireland

To temporally anchor the major historical admixture events into Ireland we used GLOBETROTTER [23] with modern surrogate populations represented by 4,514 Europeans [24] and 1,973 individuals from the PoBI dataset [7], excluding individuals sampled from Northern Ireland. Of all the European populations considered, ancestral influence in Irish genomes was best represented by modern Scandinavians and northern Europeans, with a significant single-date one-source admixture event overlapping the historical period of the Norse-Viking settlements in Ireland (p < 0.01; fit quality FQ_B_ > 0.985; Fig 6). This was recapitulated to varying degrees in specific genetically- and geographically-defined groups within Ireland, with the strongest signals in south and central Leinster (the largest recorded Viking settlement in Ireland was *Dubh linn* in present-day Dublin), followed by Connacht and north Leinster/Ulster (S5 Fig; S6 Table). This suggests a contribution of historical Viking settlement to the contemporary Irish genome and contrasts with previous estimates of Viking ancestry in Ireland based on Y chromosome haplotypes, which have been very low [25]. The modern-day paucity of Norse-Viking Y chromosome haplotypes may be a consequence of drift with the small patrilineal effective population size, or could have social origins with Norse males having less influence after their military defeat and demise as an identifiable community in the 11th century, with persistence of the autosomal signal through recombination.

**Fig 6.**
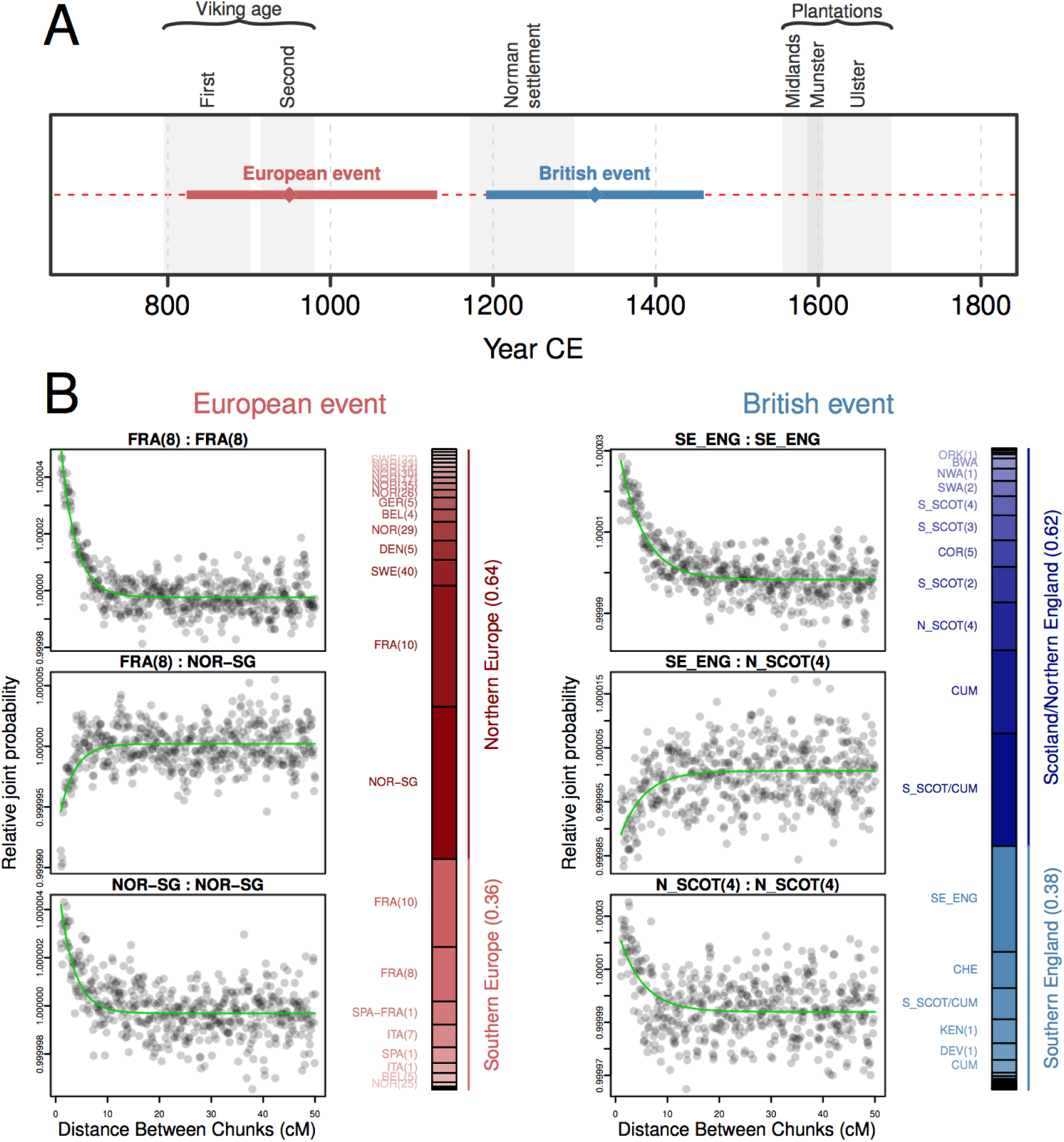
All-Ireland GLOBETROTTER admixture date estimates for European and British surrogate admixing populations. A summary of the date estimates and 95% confidence intervals for inferred admixture events into Ireland from European and British admixing sources is shown in **(A)**, with ancestry proportion estimates for each historical source population for the two events and example coancestry curves shown in **(B)**. In the coancestry curves *Relative joint probability* estimates the pairwise probability that two haplotype chunks separated by a given genetic distance come from the two modeled source populations respectively (ie FRA(8) and NOR-SG); if a single admixture event occurred, these curves are expected to decay exponentially at a rate corresponding to the number of generations since the event. The green fitted line describes this GLOBETROTTER fitted exponential decay for the coancestry curve. If the sources come from the same ancestral group the slope of this curve will be negative (as with FRA(8) vs FRA(8)), while a positive slope indicates that sources come from different admixing groups (as with FRA(8) vs NOR-SG). The adjacent bar plot shows the inferred genetic composition of the historical admixing sources modelled as a mixture of the sampled modern populations. A European admixture event was estimated by GLOBETROTTER corresponding to the historical record of the Viking age, with major contributions from sources similar to modern Scandinavians and northern Europeans and minor contributions from southern European-like sources. For admixture date estimates from British-like sources the influence of the Norman settlement and the Plantations could not be disentangled, with the point estimate date for admixture falling between these two eras and GLOBETROTTER unable to adequately resolve source and proportion details of admixture event (fit quality FQ_B_< 0.985). The relative noise of the coancestry curves reflects the uncertainty of the British event. Cluster labels (for the European clustering dendrogram, see S4 Fig; for the PoBI clustering dendrogram, see S3 Fig): FRA(8), France cluster 8; NOR-SG, Norway, with significant minor representations from Sweden and Germany; SE_ENG, southeast England; N_SCOT(4) northern Scotland cluster 4.

European admixture date estimates in northwest Ulster did not overlap the Viking age but did include the Norman period and the Plantations (S5 Fig). This may indicate limited Viking activity in Ulster, or, that due to the similarity in sources for the Viking and Anglo-Norman invasions and the Plantations, GLOBETROTTER failed to disentangle the earlier events from the later. This is not unexpected given the extent of the Plantations in Ulster [26], the relative timings of the invasions and the degree of Viking involvement in Britain and Europe. Indeed, when considering Britain as an admixing source using PoBI data, GLOBETROTTER date estimates for northwest Ulster overlapped the Plantations, although for other regions in Ireland (and for Ireland considered as a whole) these admixture events were less clearly defined, likely reflecting a history of continuous gene flow between the two islands in the prevailing years (Fig 6; S5 Fig and S7 Table). The all-Ireland point estimate for admixture from Britain spanned the Norman settlement instead of the Plantations, but GLOBETROTTER was unable to adequately resolve the model details for this event (fit quality FQ_B_ < 0.985; Fig 6), indicating that this estimate is not a good reflection of the true timings and extent of admixture from Britain. As noted in the PoBI study, the overall influence of British admixture in Ireland (and *vice versa*) has involved extensive and constant gene flow before, during and after the major population movements detailed in Fig 6, with particular swells of peopling during the Plantations. The genetic legacies of the populations of Ireland and Britain are therefore extensively intertwined and, unlike admixture from northern Europe, too complex to model with GLOBETROTTER.

### Conclusions

Our results show that population structure is detectable on the island of Ireland and is consistent with a combination of the homogenizing effect of geographically punctuated admixture and diversification among Celtic subpopulations. The inclusion of Irish data with British samples from the PoBI study provides an anchor for Celtic ancestry in the British Isles, filling out the genetic landscape of the islands. It is also clear that historical migrations into Ireland have left a greater genomic footprint than previously anticipated; our consideration of autosomal data escapes the constraints of patrilineal genetics and has allowed us to detect a much greater Viking influence than previously estimated with Y chromosome data. Although the genetic imprint of the British Plantations is much harder to delineate, the inter-island exchange and clustering observed between present-day individuals from Northern Ireland and Scotland signals the enduring impact of these historical movements of people.

Unlike the PoBI study, Irish data were not specifically selected for longstanding pure ancestry in each geographic region (for example, having four grandparents in a location), but instead represent a repurposed medical dataset. Our data are therefore more representative of those that are typically used in population-based genome-wide surveys for trait-associated genetic variation; as these studies survey increasingly rare genetic variants in larger populations, the geospatial segregation of rare haplotypes and variants will become increasingly important, especially when environmental effects and interactions play a role [27]. Our observation that these haplotypes are intricately tied to geography in Ireland and Britain highlights the importance of considering fine-grained population structure in future studies.

## Methods

### Data and quality control

Our study included three datasets of genotype data: a population-based Irish ALS case-control dataset (n = 991) incorporating existing [28] and newly-genotyped samples, the People of the British Isles dataset (EGA accession ID EGAD00010000632; n = 2,020) [7] and a pan-European dataset derived from a genome-wide association study (GWAS) for multiple sclerosis (MS; EGA accession ID EGAD00000000120; n = 4,514) [24] (S1 Text: Populations). All Irish subjects provided written informed consent to participate in genetic research and the study was approved by the Beaumont Hospital Research Ethics Committee in Dublin, Ireland. We applied quality control to each dataset using PLINK 1.9 [29] and merged data as detailed in Supplementary Methods (S1 Text: Quality Control). Briefly, we excluded both infrequent and high-missingness SNPs; individuals with high missingness, excessive heterozygosity or cryptic relationships to other individuals in the data; and finally individuals who had been removed during QC carried out in the source papers.

As the European dataset included patients and controls from a GWAS for MS, we additionally removed SNPs in a 15 Mb region surrounding the strongly associated HLA locus on chromosome 6 (GRCh37 position chr6:22,915,594-37,945,593), as is consistent with previous studies using the data [7,[30]. This was to avoid haplotypic bias arising from this association.

The final post-QC Irish (n = 991), British (n = 2,020) and European datasets (n = 4,514) contained 407,750 SNPs, 521,883 SNPs and 363,396 SNPs at zero missingness, respectively. The final merge of British and Irish data (n = 3,008) and European and Irish data (n = 5,506) contained 214,632 SNPs and 166,139 SNPs respectively at zero missingness. Further details regarding samples and QC per dataset are described in Supplementary Methods (S1 Text: Populations and S1 Text: QC)

Geographic information was available for 544 of the 991 Irish samples in the form of home address. To preserve anonymity this was jittered in all maps containing patients (Fig 1 and S5 Fig). For all British and some Northern Irish data, sample location was supplied by the authors of PoBI [7] as membership of 35 sampling regions. Finally, for European data sampling country was available [24]. Full details of treatment of samples for mapping are available in Supplementary methods (S1 Text: Mapping.)

### Phasing

We phased autosomal genotypes in each dataset and merged dataset with SHAPEIT V2 [31] using the 1000 Genomes (Phase 3) as a reference panel [32]. A pre-phasing step was carried out (--check) to remove any SNPs which did not correctly align to the 1000 genomes reference panel. Samples were then split by chromosome and phased together using default settings and the GRCh37 build genetic map to estimate linkage disequilibrium.

### fineSTRUCTURE analysis

To detect population structure we performed ChromoPainter/fineSTRUCTURE analysis [14] on each of the population datasets (Irish, British and European) individually, and then separately on a merge of the Irish and British datasets. In brief, we used ChromoPainter to paint each individual using all other individuals (-a 0 0) using default settings with the exception that the number of “chunks” per region value was set to 50 (-k 50) for all analyses including Irish and British individuals to account for the longer haplotypes observed in these datasets, in keeping with previous studies [7,[30]. The fineSTRUCTURE algorithm was then run on the resulting coancestry matrix to determine genetic clusters based on patterns of haplotype sharing. Further details are included in the Supplementary Methods (S1 Text: fineSTRUCTURE analysis).

### Cluster robustness

We assessed the robustness of Irish clusters by calculating total variation distance (TVD) as described in the PoBI study [7]. This metric compares the “copying vectors” of pair of clusters. Here we define the copying vector for a given cluster *A* as a vector of the average lengths of DNA donated by each cluster to individuals within cluster *A* under the ChromoPainter model. Hence the magnitude of differences between copying vectors of two clusters reflects the distances between those clusters in terms of their haplotypic sharing with other clusters. TVD can therefore be used to determine whether fineSTRUCTURE clusters detect significant differences in haplotype sharing, and hence ancestry.

We tested whether the observed clustering performed better than chance by permuting (1,000 times) the individuals in each of our cluster pairings into clusters of the same size, and calculating the number of permutations that exceeded our original TVD score. If 1,000 unique permutations were not possible, the maximum number of unique permutations was used instead. P-values were calculated based on the number of permutations greater than or equal to the original TVD statistic. All p-values for Irish clusters were less than or equal to 0. 001 indicating robust clustering. (S1 Table and S2 Table). We also applied these methods to our Irish cluster groups (Fig 1) and observed that these are statistically distinct (S3 Table and S4 Table).

To provide an additional measure of population differentiation between “cluster groups” we calculated mean FST between groups using PLINK 1.9 [29] which is reported in S5 Table.

### Estimating admixture dates

We used the GLOBETROTTER method [23] to infer and date admixture events from Europe and Britain into Ireland separately. GLOBETROTTER uses output from ChromoPainter to estimate the pairwise likelihood of being painted by any two surrogate populations at a variety of genetic distances to generate coancestry curves. Assuming a single admixture event, these curves are expected to follow an exponential decay rate equal to the time in generations since admixture occurred [23]. As the true admixing sources are modelled as a linear mixture of surrogate sources rather than individual sources this method has the advantage of not requiring exactly sampled source populations.

For our analysis we ran GLOBETROTTER with default settings twice to detect simple admixture into the island of Ireland as a whole, as well as into individual genetic clusters from the Republic of Ireland (S5 Fig). European clusters (S4 Fig) and British clusters (S3 Fig) were used as surrogate populations to represent the admixing sources in two independent analyses. Target and donor clusters for this analysis were defined using the fineSTRUCTURE maximum concordance tree method described in PoBI ([7]) to ensure homogeneity (Supplementary methods S1 Text: fineSTRUCTURE analysis); hence, the Irish target clusters that were used differ slightly from those in Fig 1. Briefly, for each surrogate population separately (Europe and Britain) we applied ChromoPainter v2 to paint Ireland and the surrogate population using the surrogate population as donors and generated a copying matrix (chunk lengths) for all individuals, and also 10 painting samples for each Irish individual as recommended. GLOBETROTTER was then run for 5 mixing iterations twice, first using the null.ind:1 setting to test for any evidence of admixture and then null.ind:0 setting to infer dates and sources. We ran 100 bootstraps for admixture date and calculated the probability of a null model of no admixture as the proportion of nonsensical inferred dates (<1 or >400 generations) produced by the null.ind:1 model, as in the GLOBETROTTER study [23]. Confidence intervals for the date were calculated from the bootstraps for the standard model (null.ind: 0) using the empirical bootstrap method. (See S1 Text: Globetrotter analysis of Admixture Dates for further details). A generation time of 28 years was assumed as in previous studies of this nature [7,[23] for conversion of all date estimates from generations to years.

### Ancestry proportion estimation

We assessed the ancestral make up of Ireland in terms of Europe and Britain for each Republic of Ireland cluster (see Estimating admixture dates) to explore variation in ancestry across Ireland. To do so we modelled each cluster’s average genome as a linear mixture of the European and British donor populations using the method described in the PoBI study [7] and implemented in GLOBETROTTER (num.mixing.iterations: 0). This approach uses the ChromoPainter chunk length output to estimate the proportion of DNA which most closely coalesces with each individual from the donor populations, correcting for noise caused by similarities between donor populations whose splits may have occurred after the coalescence event. This is achieved through a multiple linear regression of the form Y_p_ = B_1_X_1_ + B_2_X_2_ + … +B_g_X_g_, where Y_p_ is a vector of the averaged length (cM) of DNA that individuals across cluster P copy from each donor group, normalised to sum to 1 across all donor groups, and X_g_ is the vector describing the average proportion of DNA that individuals in donor group g copy from other donor groups including their own. The coefficients of this equation B_1_…B_g_ are thus interpreted as the “cleaned” proportions of the genome ancestral to each donor group. The equation is solved using a non-negative-least squares function such that B_g_ ≥ 0 and the sum of proportions across groups equals 1.

To assess uncertainty of these ancestry proportion estimates we again follow PoBI [7] and resample from the ChromoPainter chunk length output to generate N_p_ pseudo individuals for each cluster P. We achieve this by randomly sampling each of the autosomal chromosome pairs 1-22 with replacement N_p_ times from the pool of all autosomal chromosomes pairs 1-22 across all individuals within that cluster, and then randomly summing sets of 22 of these chromosome pairs to generate each pseudo individual. We then use these N_p_ pseudo individuals as a bootstrap for Y_p_ above and solve for B_g_. We resampled 1,000 times per cluster and used the inner 95% quantiles of this sampling distribution to estimate confidence intervals for the sample.

For comparison we implemented an alternative delete one chromosome jack-knife approach as in Montinaro et al. [33.], and estimated the s.e. as in ref. [34] (S6 Fig and S7 Fig.)

We also used this linear regression model to determine per-individual ancestry proportion estimates from different British clusters across Ireland, treating each individual as a cluster to enable us to assess whether gene flow from northern Britain had a gradient across Ireland.

### ADMIXTURE

To estimate the proportion of British admixture into Irish clusters, ADMIXTURE [18] was run on the combined PoBI and Irish datasets, alongside eighteen ancient individuals from the Iron Age, Roman and Anglo-Saxon periods of northeast and southeast England [16,[17]. Pseudo-haploid genotypes were generated for the ancient genomes at the relevant variant sites, as is standard for low coverage data, and subsequently merged with the modern diploid dataset. Data were then pruned for linkage disequilibrium between SNPs using PLINK 1.9 (r^2^ > 0.25 in a sliding window of 1000 SNPs advancing 50 SNPs each time) resulting in 86,481 remaining SNPs. No missingness was allowed for modern individuals, with a range of 33,643-85,553 sites used for ancient samples. Following ADMIXTURE estimation, cross-validation error was calculated using the --cv flag for 5 iterations to determine the K value for which the model has the best predictive accuracy (K=2). Additionally 200 bootstraps of the data were run to estimate the standard error of the parameters using the −B flag. This British admixture component was regressed against PC2 of the Irish ChromoPainter coancestry matrix to determine the role of British ancestry in the differentiation of PC2 in Ireland. We also performed analysis of variance (ANOVA) on British admixture component per cluster group to identify if cluster by cluster differences existed.

### PCA and t-SNE

ChromoPainter coancestry matrices were projected in lower-dimensional space using principal component analysis (PCA) and t-distributed stochastic neighbour embedding (t-SNE) [21]. PCA was run using the default approach provided as part of the fineSTRUCTURE R tools [14] (http://www.paintmychromosomes.com). The R package Rtsne (https://github.com/ikrijthe/Rtsne) was used to construct a 2-dimensional embedding of the ChromoPainter coancestry matrix over 5,000 iterations using a perplexity of 30, a learning rate of 200 and an initial PCA calculated over 100 dimensions. Several t-SNE runs were performed to assess concordance between embedding solutions.

### Other statistical analyses

All linear regressions and ANOVA tests were carried out in base statistics package in R version 3.2.3 [35].

## Acknowledgements

We gratefully acknowledge Dr T. Rowan McLaughlin for providing helpful insights and critical review of the manuscript. A subset of the Irish data was generated as part of Project MinE (www.projectmine.com) and we acknowledge the DJEI/DES/SFI/HEA Irish Centre for High-End Computing (ICHEC) for the provision of computational facilities and support.

## References

1. Bayliss A, Woodman P. A New Bayesian Chronology for Mesolithic Occupation at Mount Sandel, Northern Ireland. Proceedings of the Prehistoric Society. 2009;75: 101–123.

2. Cassidy LM, Martiniano R, Murphy EM, Teasdale MD, Mallory J, Hartwell B, et al. Neolithic and Bronze Age migration to Ireland and establishment of the insular Atlantic genome. Proc Natl Acad Sci U S A. 2016;113: 368–373.

3. Dillon M, Chadwick NK. The Celtic Realms. Weidenfeld & Nicolson; 1967.

4. Jones C. The Edinburgh History of the Scots Language. Edinburgh University Press; 1997.

5. Koch JT. Celtic culture: a historical encyclopedia. ABC-CLIO; 2006.

6. Duffy S. The concise history of Ireland. Gill & Macmillan; 2000.

7. Leslie S, Winney B, Hellenthal G, Davison D, Boumertit A, Day T, et al. The fine-scale genetic structure of the British population. Nature. 2015;519: 309–314.

8. Haak W, Lazaridis I, Patterson N, Rohland N, Mallick S, Llamas B, et al. Massive migration from the steppe was a source for Indo-European languages in Europe. Nature. 2015;522: 207–211.

9. Lazaridis I, Nadel D, Rollefson G, Merrett DC, Rohland N, Mallick S, et al. Genomic insights into the origin of farming in the ancient Near East. Nature. 2016;536: 419–424.

10. O’Dushlaine CT, Morris D, Moskvina V, Kirov G, International Schizophrenia Consortium, Gill M, et al. Population structure and genome-wide patterns of variation in Ireland and Britain. Eur J Hum Genet. 2010;18: 1248–1254.

11. McLaughlin RL, Kenna KP, Vajda A, Heverin M, Byrne S, Donaghy CG, et al. Homozygosity mapping in an Irish ALS case-control cohort describes local demographic phenomena and points towards potential recessive risk loci. Genomics. 2015;105: 237–241.

12. Moore LT, McEvoy B, Cape E, Simms K, Bradley DG. A Y-chromosome signature of hegemony in Gaelic Ireland. Am J Hum Genet. 2006;78: 334–338.

13. McEvoy B, Simms K, Bradley DG. Genetic investigation of the patrilineal kinship structure of early medieval Ireland. Am J Phys Anthropol. Wiley Subscription Services, Inc., A Wiley Company; 2008;136: 415–422.

14. Lawson DJ, Hellenthal G, Myers S, Falush D. Inference of population structure using dense haplotype data. PLoS Genet. 2012;8: e1002453.

15. Duffy S. Atlas of Irish History. Gill & MacMillan; 2012.

16. Martiniano R, Caffell A, Holst M, Hunter-Mann K, Montgomery J, Müldner G, et al. Genomic signals of migration and continuity in Britain before the Anglo-Saxons. Nat Commun. 2016;7: 10326.

17. Schiffels S, Haak W, Paajanen P, Llamas B, Popescu E, Loe L, et al. Iron Age and Anglo-Saxon genomes from East England reveal British migration history. Nat Commun. 2016;7: 10408.

18. Alexander DH, Novembre J, Lange K. Fast model-based estimation of ancestry in unrelated individuals. Genome Res. 2009;19: 1655–1664.

19. Commission E-E, Others. Regions in the European Union. Nomenclature of territorial units for statistics. NUTS 2010/EU-27. Luxemburgo: Publications Office of the European Union; 2011.

20. Novembre J, Johnson T, Bryc K, Kutalik Z, Boyko AR, Auton A, et al. Genes mirror geography within Europe. Nature. 2008;456: 98–101.

21. van der Maaten L. Learning a Parametric Embedding by Preserving Local Structure. Proceedings of Machine Learning Research. 2009;5: 384–391.

22. Deacon B. A Concise History of Cornwall. University of Wales Press-Hi; 2007.

23. Hellenthal G, Busby GBJ, Band G, Wilson JF, Capelli C, Falush D, et al. A genetic atlas of human admixture history. Science. 2014;343: 747–751.

24. International Multiple Sclerosis Genetics Consortium, Wellcome Trust Case Control Consortium 2, Sawcer S, Hellenthal G, Pirinen M, Spencer CCA, et al. Genetic risk and a primary role for cell-mediated immune mechanisms in multiple sclerosis. Nature. 2011;476: 214–219.

25. McEvoy B, Brady C, Moore LT, Bradley DG. The scale and nature of Viking settlement in Ireland from Y-chromosome admixture analysis. Eur J Hum Genet. 2006;14: 1288–1294.

26. McLaughlin R, Lyttleton J. An Archaeology of Northern Ireland, 1600-1650. Department for Communities; 2017.

27. Mathieson I, McVean G. Differential confounding of rare and common variants in spatially structured populations. Nat Genet. 2012;44: 243–246.

28. van Rheenen W, Shatunov A, Dekker AM, McLaughlin RL, Diekstra FP, Pulit SL, et al. Genome-wide association analyses identify new risk variants and the genetic architecture of amyotrophic lateral sclerosis. Nat Genet. 2016;48: 1043–1048.

29. Chang CC, Chow CC, Tellier LC, Vattikuti S, Purcell SM, Lee JJ. Second-generation PLINK: rising to the challenge of larger and richer datasets. Gigascience. 2015;4: 7.

30. Gilbert E, Carmi S, Ennis S, Wilson JF, Cavalleri GL. Genomic insights into the population structure and history of the Irish Travellers. Sci Rep. 2017;7: 42187.

31. Delaneau O, Marchini J, Zagury J-F. A linear complexity phasing method for thousands of genomes. Nat Methods. 2011;9: 179–181.

32. Auton A, Abecasis GR, Altshuler DM, Durbin RM, Abecasis GR, Bentley DR, et al. A global reference for human genetic variation. Nature. 2015;526: 68–74.

33. Montinaro F, Busby GBJ, Pascali VL, Myers S, Hellenthal G, Capelli C. Unravelling the hidden ancestry of American admixed populations. Nat Commun. 2015;6: 6596.

34. Frank M T, Meijer E, Van Der Leeden R. Delete-m Jackknife for Unequal m. Stat Comput. Kluwer Academic Publishers; 1999;9: 3–8.

35. CoreTeam R. R: A Language and Environment for Statistical Computing. Vienna, Austria: R Foundation for Statistical Computing; 2015. 2015.

